# A preclinical model to investigate normal tissue damage following fractionated radiotherapy to the head and neck

**DOI:** 10.1101/2022.05.19.492439

**Authors:** Inga Solgård Juvkam, Olga Zlygosteva, Delmon Arous, Hilde Kanli Galtung, Eirik Malinen, Tine Merete Søland, Nina Jeppesen Edin

## Abstract

Radiotherapy of head and neck cancer is known to cause both early and late-occurring toxicities. To better appraise normal tissue responses and their dependence on treatment parameters such as radiation field and type, as well as dose and fractionation scheme, a preclinical model with relevant endpoints is required. 12-week old female C57BL/6J mice were irradiated with 100 or 180 kV X-rays to total doses ranging from 30 to 85 Gy, given in 10 fractions over 5 days. The radiation field covered the oral cavity, swallowing structures, and salivary glands. Monte Carlo simulations were employed to estimate tissue dose distribution. The follow-up period was 35 days, in order to study the early radiation-induced effects. Baseline and post irradiation investigations included macroscopic and microscopic examinations of the skin, lips, salivary glands, and oral mucosa. Saliva sampling was performed to assess the salivary gland function following radiation exposure. A dose dependent radiation dermatitis in the skin was observed for doses above 30 Gy. Oral mucositis in the tongue appeared as ulcerations on the ventral surface of the tongue for doses of 75-85 Gy. The irradiated mice showed significantly reduced saliva production compared to controls. In summary, a preclinical model to investigate a broad panel of normal tissue responses following fractionated irradiation of the head and neck region was established. The optimal dose to study early radiation-induced effects was found to be around 75 Gy, as this was the highest tolerated dose that gave acute effects similar to what is observed in cancer patients.

## Introduction

Head and neck (H&N) cancer patients who receive radiotherapy (RT) as part of their treatment may be severely affected by radiation-induced damages to normal tissue. RT can result in early side effects such as dermatitis and oral mucositis, which may occur during or soon after RT and could potentially lead to interruption of the treatment. Furthermore, RT can also produce late side effects that may severely reduce the patient’s quality of life, such as salivary gland hypofunction, tissue fibrosis, and osteoradionecrosis (1, 2). Generally, clinical symptoms of side effects following RT are well documented. However, preclinical models are essential to investigate radiation-induced early and late side effects in normal tissues of the H&N region and their dependence on RT-related factors such as radiation field and type (e.g., X-rays and protons) as well as dose and fractionation scheme. A better appraisal of these effects and underlying biological processes may aid the development of new methods and strategies for mitigating or eliminating such tissue damage and subsequent clinical symptoms (3).

Current radical RT of H&N cancer is delivered as fractionated treatment, normally in fractions of 2 Gy to a total dose of typically 70 Gy (4). Still, most previous studies investigating the side effects of RT in the H&N region of murine model systems have used either single dose (SD) irradiation or extremely hypofractionated schedules with few fractions of very high doses (5-10). Some studies have used fractionation for H&N experiments (11-14), but without any justification for the choice of dose. Also, the radiation field(s) employed in previous studies did not cover the typical region as seen in patients, which should encompass areas such as the oral cavity, the oropharynx, and the laryngopharynx. Examples from the literature encompass large radiation fields including the entire head or upper body of the animal (5, 8, 10) or small radiation fields only covering e.g., the tongue or snout of the animal (9, 11-13), which differ from radiation fields used in the clinic. Thus, preclinical studies employing more clinically relevant radiation fields and fractionated radiation delivery are highly needed. Also, as a broad spectrum of tissue responses may occur after irradiation of the H&N area, the selected preclinical model should be an optimal compromise with respect to ease of use and possibilities for long term follow-up with tissue and liquid sampling.

The aim of the present investigation was to establish a preclinical model employing a clinically relevant radiation field and careful assessment of radiation dose distribution to study various early radiation-induced effects in the H&N region induced by different dose deliveries.

## Materials and Methods

### Animals

Nine-week-old C57BL/6J male and female mice were purchased from Janvier (France), kept in a 12-h light/12-h dark cycle under pathogen-free conditions and fed a standard commercial fodder with water given *ad libitum*. Standard housing with nesting material and refuge was provided. All experiments were approved by the Norwegian Food Safety Authority (ID 20889, 26246 and 27931) and performed in accordance with directive 2010/63/EU on the protection of animals used for scientific purposes. At the onset of experiments, animals were 12 weeks old.

### Irradiation Procedure

Radiotherapy was delivered in 10 fractions over 5 days (8 am and 4 pm) with a Faxitron Multirad225 irradiation system (Faxitron Bioptics, Tucson, AZ, USA). Four sets of experiments were performed with two different X-ray settings: (1) 180 kV X-ray potential, 10 mA current, 0.3 mm Cu filter, and 0.65 Gy/min dose rate, and (2) 100 kV, 15 mA, 2.0 mm Al, and 0.75 Gy/min. Absolute calibration of the X-ray system was conducted using an FC65-G ionization chamber (IBA Dosimetry, Germany) together with a MAX-4000 electrometer (Standard Imaging, USA) according to standards for absorbed dose to water. For all fractions, animals were anesthetized using gas anesthesia with Sevoflurane 4 % in O_2_. The anesthetized mice were positioned on their right side in a custom-made foam holder with the beam entering on the left side. A lead collimator was custom built to define a radiation field of 25 × 20 mm covering the oral cavity, pharynx, and major salivary glands and placed on top of the foam holder. The radiation field was carefully planned to only irradiate the tissues of interest and avoid exposure of the eyes and brain. A built-in X-ray imaging system was used to verify the anatomical location of the radiation field (Fig. 1A). In the following, day 0 is the time point where the first irradiation was performed.

**Figure 1.**
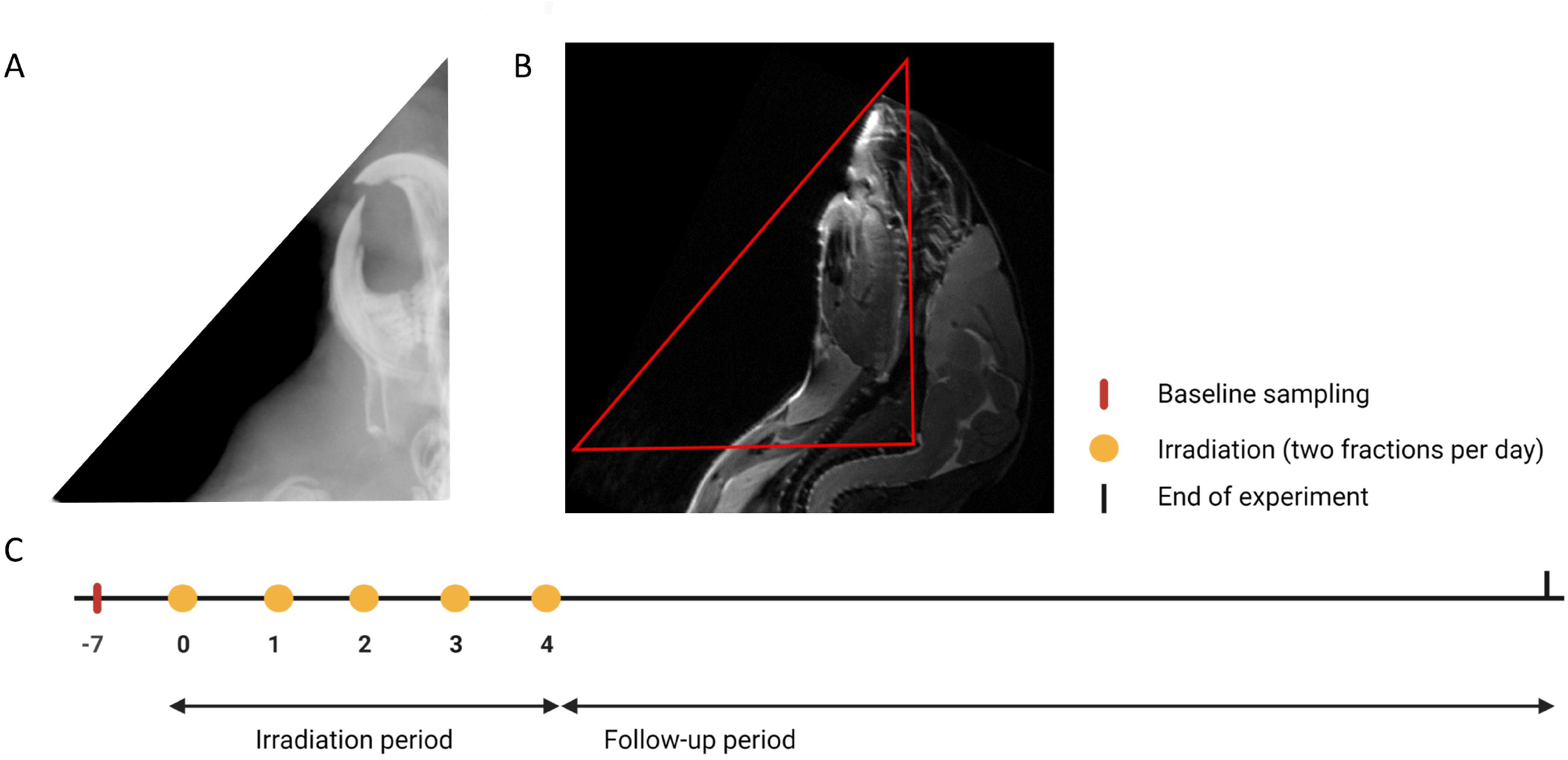
Experimental design. (A) X-ray image of the radiation field used. (B) MR image of the head and neck region. The red triangle shows the approximate radiation field. (C) Experimental protocol timeline. Irradiation was given twice a day for 5 days (days 0-4) as 10 fractions of 3, 4.4, 5, 5.75, 6.5, 7.5 and 8.5 Gy. Examinations of the oral cavity and saliva sampling were performed throughout the follow-up period.

### MRI

Magnetic Resonance Imaging of the H&N region was performed using a 7.05 T Biospec scanner (Bruker Medical systems, Germany) on the same days as the saliva sampling. A fast T2 weighted spin-echo sequence, TurboRARE, with TE = 31 ms and TR = 3100 ms was employed. Body temperature was monitored and maintained at 37 °C by a feedback-regulated heating fan. Respiration rate was monitored by a respiration probe. The MR image was used to show the radiation field in relation to the anatomical structures of the mouse (Fig. 1B). For MRI, animals were anesthetized using gas anesthesia with Sevoflurane 4 % in O_2_.

### Monte Carlo Simulations

Monte Carlo (MC) simulations of the dose distribution in mice were conducted in FLUKA 4-1.1 (15) (see Supplementary file for details). Briefly, the simulations were performed in computed tomography (CT) images of one euthanized male mouse (10-11 weeks of age). Both X-ray setting 1 and 2 were simulated, providing 3D dose distributions for 100 kV and 180 kV X-ray spectra, respectively. A rectangular irradiation field was used which covered the same regions as defined experimentally. The transport and production cutoff of photons and electrons was set to 1 keV. The treatment field was simulated for 5 × 10^7^ primary X-ray photons. The absorbed dose was scored on a voxel-by-voxel basis, providing tissue-specific dose estimates where mean and ranges are reported.

### Experimental Protocol

Altogether, four sets of experiments were accomplished. First, two pilot experiments (experiment 1 and 2) were performed to establish procedures and appropriate X-ray voltages and doses. Then, two main experiments (experiment 3 and 4) were conducted with a larger number of animals in each treatment group to further assess the early radiation-induced effects. Experiment 1 employed both male and female mice, randomly assigned to either sham treatment, 10 × 3 Gy, or 10 × 4.4 Gy irradiation (n = 3 for each gender and dose group). Here, X-ray setting 1 (see *Irradiation Procedure* above) was used. An aggressive behavior was observed in the male group that led to loss of several mice during experiment 1. Because possible gender difference in response to radiation was not the aim of the current study, we decided to include only female mice in the further experiments. Therefore, experiment 2 used only female mice, randomly assigned to either sham treatment, 10 × 5 Gy, 10 × 5.75 Gy, or 10 × 6.5 Gy irradiation (n = 4 for each treatment group) with a shorter follow-up period to ensure documentation of histological effects. As will become evident, pilot experiment 1 and 2 gave rather mild symptoms and experiment 3 was therefore conducted with increased doses. Experiment 3 used only female mice, randomly assigned to either sham treatment or 10 × 8.5 Gy (n = 10 for each treatment group). However, the dose of 10 × 8.5 Gy was not well tolerated by the mice and experiment 3 was terminated on day 14 due to unacceptable weight loss (>20%). Based on experiments 1-3, the optimal dose was determined to be between 6.5 and 8.5 Gy per fraction (Gy/f). Therefore, experiment 4 was conducted with female mice, randomly assigned to either sham treatment or 10 × 7.5 Gy (n = 10 for each treatment group). In experiment 2-4, X-ray setting 2 was used because of a more clinically relevant dose distribution (presented in Results). All reported tissue doses are mean doses at the midpoint in the X-ray path through the mouse.

On day -7, baseline body weight was determined and saliva sampling was performed in all animals. On days 0-4, fractionated irradiation was given twice a day to the irradiation groups, as explained above (see also Fig.1C). The maximum follow-up period in this study was 35 days to fully cover the early radiation-induced effect. During the follow-up period, the animals were monitored frequently. Examinations of the oral cavity were performed every second day and weighing was done daily. Macroscopic examinations of the oral cavity were performed using magnifying glasses (3.5x) and a light source, while the mice were under anesthesia. Subcutaneous injection of anesthesia was used (Zoletil-mix: Zoletil-mix: 10 ml Narcoxyl or Rompun® (xylasin 20 mg/ml) + 0,5 ml Torbugesic® (butorphanol 10 mg/ml) + Zoletil® (zolazepam 125 mg and tiletamin 125 mg)). Assessment of overall activity and body weight (less than 20 % body weight loss was considered acceptable) were also performed. Presence of oral mucositis was monitored and scored as present/not present, while skin toxicity was graded using a modified scoring scheme based on the Radiation Therapy Oncology Group (RTOG) developed scoring criteria (16-18). Euthanasia was performed through overdose of anesthetic (Pentobarbitol, Exagon® Vet) by intraperitoneal injection under terminal anesthesia to prevent tissue damage from cervical dislocation.

### Histological Evaluations

At the time of euthanasia, the tongue, mucosa and skin of the lower lips, left and right buccal mucosa, and the major salivary glands (parotid, submandibular and sublingual glands) were collected and fixated for 24 hours in 10 % formalin, dehydrated in ethanol and embedded in paraffin. Tissue sections of 4 µm were cut (Leica RM2155 microtome) and stained with hematoxylin and eosin (HE) and various antibodies by immunohistochemical method (see Supplementary material). Histological examinations of the oral tissues were performed in a Nikon E90i microscope, and histological images were acquired using a Nikon DS-Ri1 camera with a CFI Plan Fluor 10x (NA 0.30), 20x (NA 0.50) or 40x objective (NA 0.75).

### Saliva Sampling

Saliva collection was performed before RT (day -7), immediately after (day 5), and at a later time point during the follow-up period (day 35). Mice were anesthetized with subcutaneous injection of Zoletil-mix (see *Experimental Protocol above*). The saliva sampling procedure was performed as previously described (19). Briefly, 0.375 mg/kg of pilocarpine (Pilocarpine hydrochloride, Sigma) was intraperitoneally administered to the mice under anesthesia. Saliva was collected into a cotton swab for 15 minutes, which was then centrifuged at 7500 g and 4 ºC for 2 minutes, and the obtained volume was measured and stored at -80 ºC. Saliva production was calculated as saliva volume (µl) per saliva collection time (15 min).

### Statistical Analyses

Statistical analyses were performed in Prism 8 for Windows (Version 8.3.0, GraphPad Software, LLC) and in RStudio (Version 4.1.1). Correlation between body weight loss and oral mucositis was analyzed using a linear regression model. Saliva production was analyzed using a mixed-effects analysis and Sidak’s multiple comparison test. Asignificance level of 0.05 was used for all comparisons.

## Results

### Monte Carlo Simulations

Figure 2 shows normalized dose maps for 100 and 180 kV X-ray voltages superimposed onto the same CT slice, which is central to the beam axis. The dose distribution is almost constant throughout the animal for 180 kV X-rays, whereas 100 kV X-rays give a gradual decline in dose. For 180 kV, the oral cavity, lip, skin and submandibular gland received on average (range) 102 (91, 111) %, 85 (73, 98) %, 80 (74, 86) % and 79 (74, 84) % of the prescribed dose, respectively. For 100 kV, the same structures received an average (range) of 93 (75, 108) %, 93 (79, 101) %, 71 (61, 79) % and 70 (64, 74) %, respectively.

**Figure 2.**
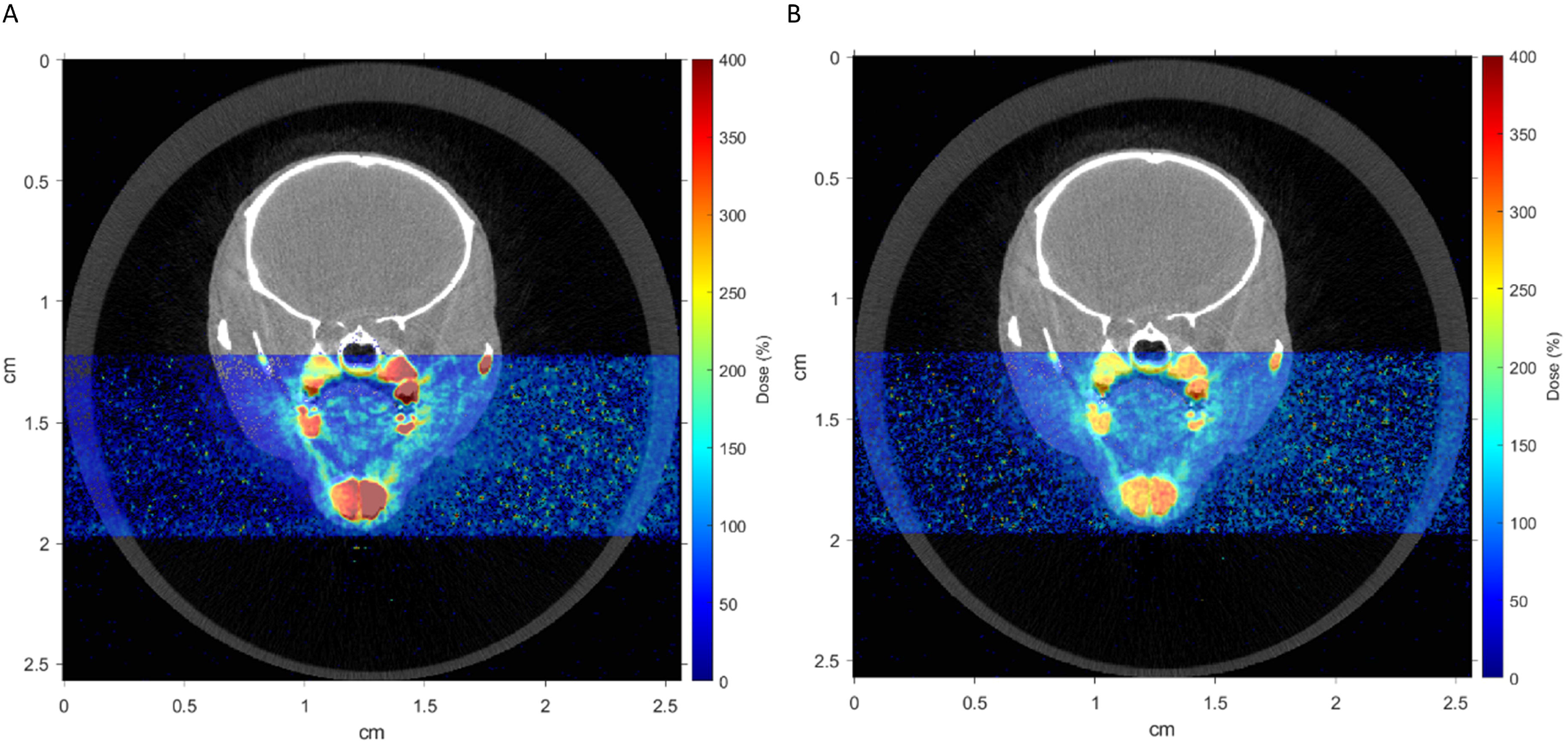
Monte Carlo stimulated dose distributions relative to the prescribed dose for (A) 100 kV and (B) 180 kV in a transverse CT image of a mouse.

### Macroscopic Changes

Early radiation-induced effects were observed macroscopically in the lower lip, the ventral part of the tongue and on the skin of the upper chest. Radiation dermatitis of the lower lip was observed in all mice exposed to more than 3 Gy/f and was graded using RTOG-based score schemes (Fig. 3A-B). Mice exposed to 4.4 Gy/f developed mild radiation dermatitis with faint erythema, mild edema, and dry desquamation, while mice exposed to 5-6.5 Gy/f developed moderate radiation dermatitis with bright erythema, moderate edema, and patchy moist desquamation. Mice exposed to 7.5 Gy/f experienced severe radiation dermatitis with confluent moist desquamation that not only affected the skin of the lower lip as in the other RT groups, but also (affected) the skin of the upper chest. Radiation dermatitis of the lower lip was first observed on day 12 and peaked about day 21 (Fig. 3C-D). LD_50_ for skin toxicity grade 1, 2 and 3 was 4.4 Gy/f, 4.7 Gy/f and 7.5 Gy/f, respectively. Skin toxicity was not fully assessed in mice exposed to 8.5 Gy/f as this group was terminated before reaching the peak of skin toxicity. Fur loss localized to the upper chest inside the radiation field was observed in all irradiation groups receiving 4.4 Gy/f or more. Areas with complete fur loss was observed typically from day 17.

**Figure 3.**
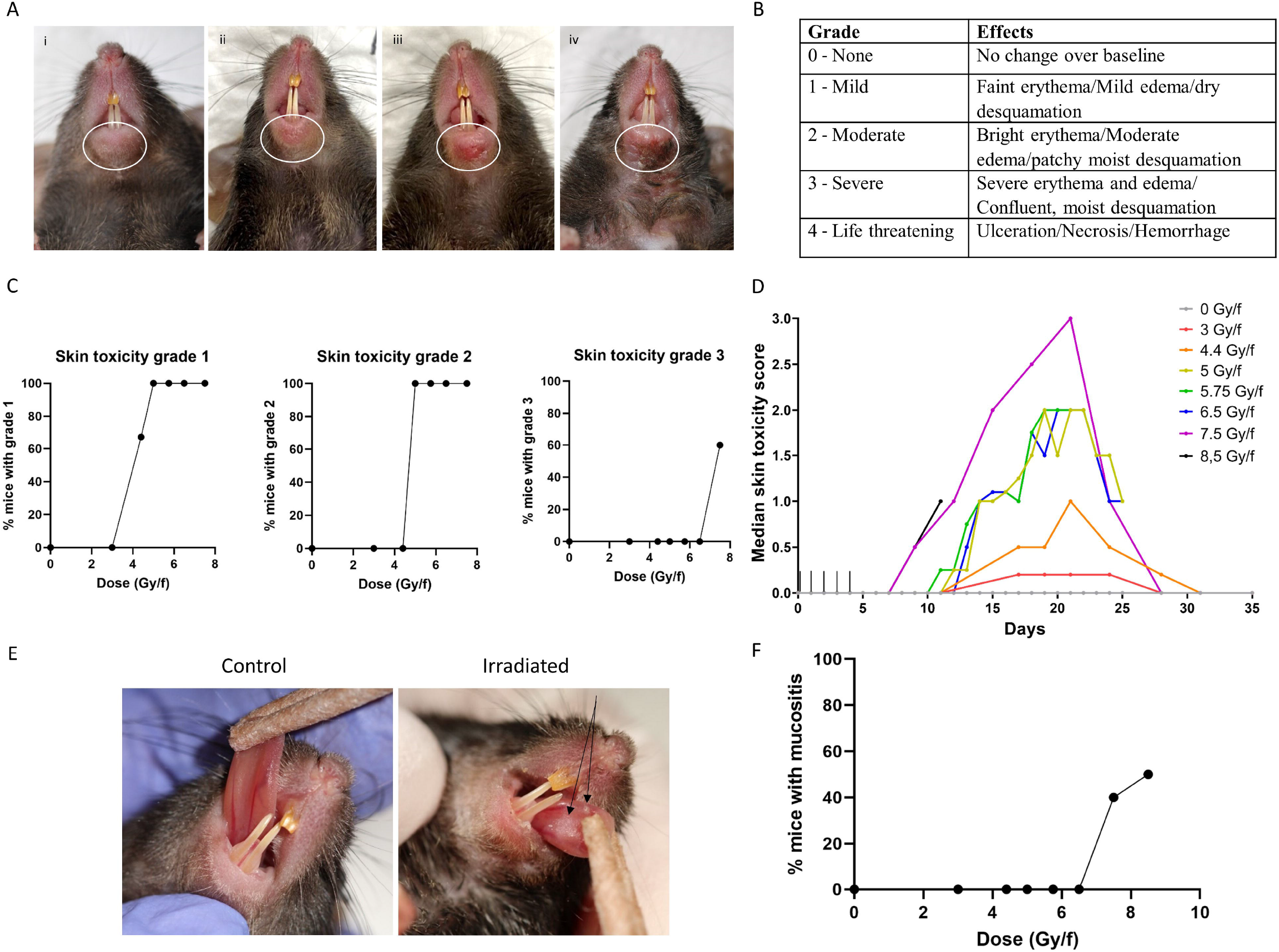
Skin toxicity effects and oral mucositis observed after fractionated irradiation with different doses per fraction. (A) Example images of different skin toxicity grades observed in the lower lip of irradiated mice; (i) grade 0, (ii) grade 1 (iii) grade 2, and (iv) grade 3. White circles show the area of interest. (B) Skin toxicity scoring scheme based on RTOG. (C) Dose response curves for different skin toxicity grades observed in different doses per fraction. (D) Timeline of median skin toxicity score observed in the skin of the lower lip of irradiated mice for different doses per fraction. The number of animals used in each group varied as following 0 Gy/f (n = 9), 3 Gy/f (n = 3), 4.4 Gy/f (n = 3), 5 Gy/f (n = 4), 5.75 Gy/f (n = 4), 6.5 Gy/f (n = 4), 7.5 Gy/f (n = 10), 8.5 Gy/f (n = 10). (E) Example images of macroscopically visible mucositis (black arrows) on the ventral tongue of mice exposed to 7.5 and 8.5 Gy/f compared to control. (F) Dose response curves for oral mucositis observed in different doses per fraction.

Oral mucositis was observed in the ventral tongue only in mice exposed to 7.5 and 8.5 Gy/f (Fig. 3E-F), and was first observed on day 9 and 12, respectively. By day 18 oral mucositis was completely resolved in the mice exposed to 7.5 Gy/f. Weight loss above 10 % was only observed in RT groups exposed to 7.5 and 8.5 Gy/f (Suppl. Fig. 1). Weight loss was significantly correlated with development of oral mucositis (p = 0.007). Mice exposed to 8.5 Gy/f experienced more than 20 % weight loss during the first 14 days of the experiment. Analgesia treatment (Temgesic injections) did not help, thus, this group was terminated at an earlier time point than planned (day 14), and 8.5 Gy/f was considered too high for this mouse strain. By termination day 14, oral mucositis was still present on the ventral tongue of the mice exposed to 8.5 Gy/f. For the present fractionation scheme and irradiation set-up, 7.5 Gy/f was thus found to be the optimal dose.

### Histological Changes

Early radiation-induced effects were also present in histologically examined tissues. Large morphological differences were seen in the lower lip between irradiated and control mice (Fig. 4A). Increased keratin thickness of the epidermis and desquamation of keratin in addition to atrophy of sebaceous glands and hair follicles was observed in the irradiated mice (Fig. 4A). Moreover, an increased amount of cells, including neutrophils, Vimentin^+^-cells and F4/80^+^-cells were seen in the connective tissue both in the skin of the lip and in the oral mucosa (Fig. 4B and Suppl. Fig. 3). Fibroblasts as well as neutrophils and lymphocytes are positive for Vimentin while macrophages in mice are F4/80^+^ (Suppl. Fig. 3). Macroscopically observed oral mucositis on the ventral tongue of mice exposed to 7.5 and 8.5 Gy/f coincided with histological findings of mucosal ulcerations (Fig. 4C). Additionally, histological examinations of the parotid glands showed acinar vacuolization on day 14 in 4 of 10 mice exposed to 8.5 Gy/f (Fig. 4D). In the submandibular and sublingual glands however, no histological differences were seen on day 14 (not shown).

**Figure 4.**
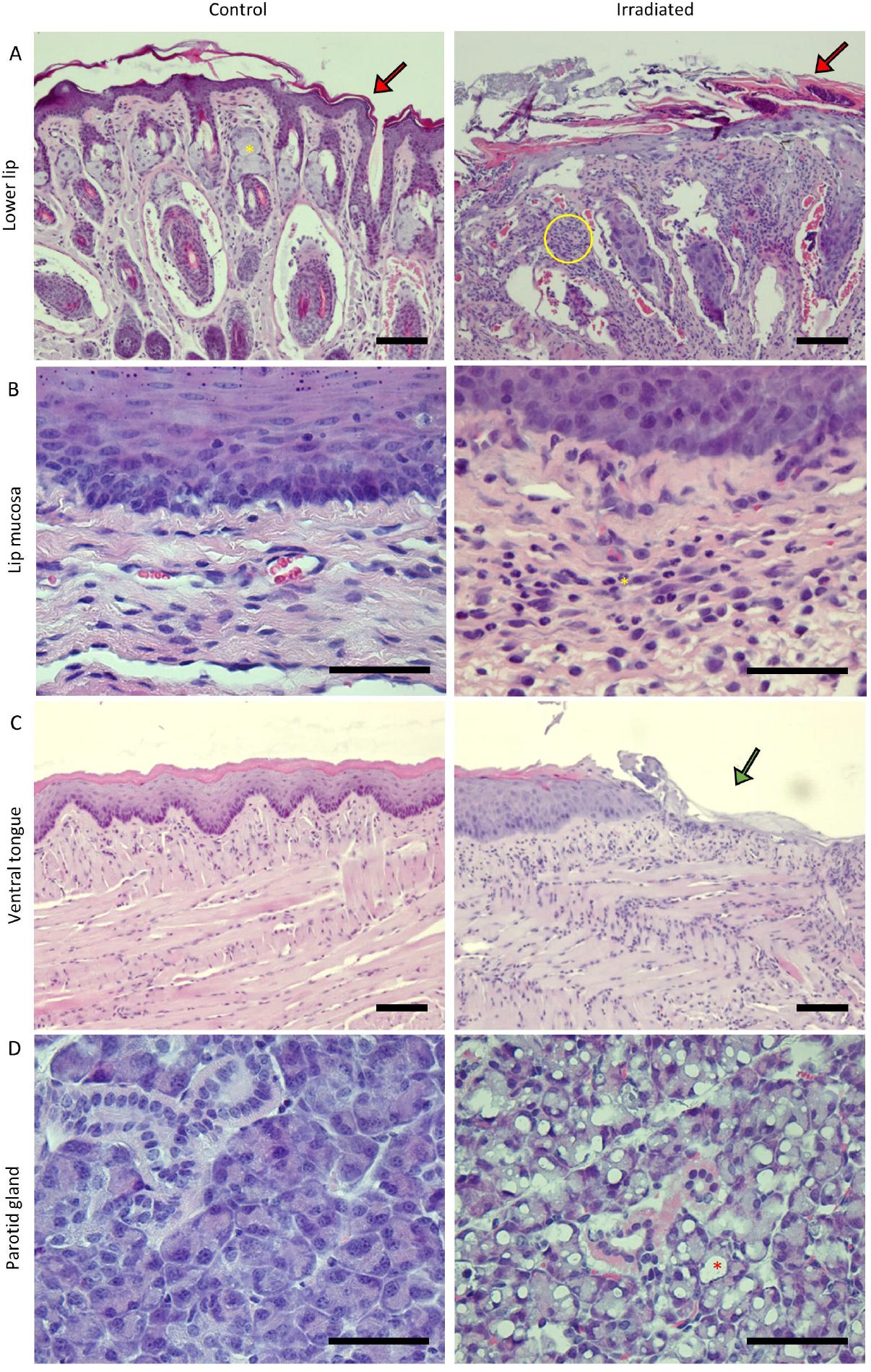
Hematoxylin and eosin (HE) stained sections of oral tissues in mice after exposure to fractionated irradiation. (A) HE sections of skin of the lower lip showed increased keratin layer (red arrow), atrophy of sebaceous glands (yellow asterisk = presence of sebaceous gland in control) and increased amount of cells in connective tissue (yellow circle) in irradiated (8.5 Gy/f on day 14) compared to control mice. Scale bar is 100 µm. (B) HE sections of lip mucosa showed increased amount of cells in connective tissue, including neutrophils (yellow asterisk), in irradiated (8.5 Gy/f on day 14) compared to control mice. Scale bar is 50 µm. (C) HE sections of the ventral tongue showing epithelial ulceration (green arrow) of the mucosa of irradiated (8.5 Gy/f on day 14) compared to control mice. Scale bar is 100 µm. (D) HE sections of the parotid gland showed acinar vacuolization (red asterisk) in irradiated (8.5 Gy/f on day 14) compared to control mice. Scale bar is 50 µm.

### Saliva production

Significantly reduced saliva production (p < 0.0001) was found in mice exposed to 7.5 Gy/f (3.6 ± 1.4 µL/15 min) compared to controls (13.1 ± 1.9 µL/15 min) on day 35 (Fig. 5). Compared to controls, there was a tendency towards reduced saliva production in the other RT groups as well (3-6.5 Gy/f), albeit not statistically significant due to large inter-individual variations and a somewhat small number of animals per group (Suppl. Fig. 4). The average coefficient of variation (CV) for saliva production was 59 %.

**Figure 5:**
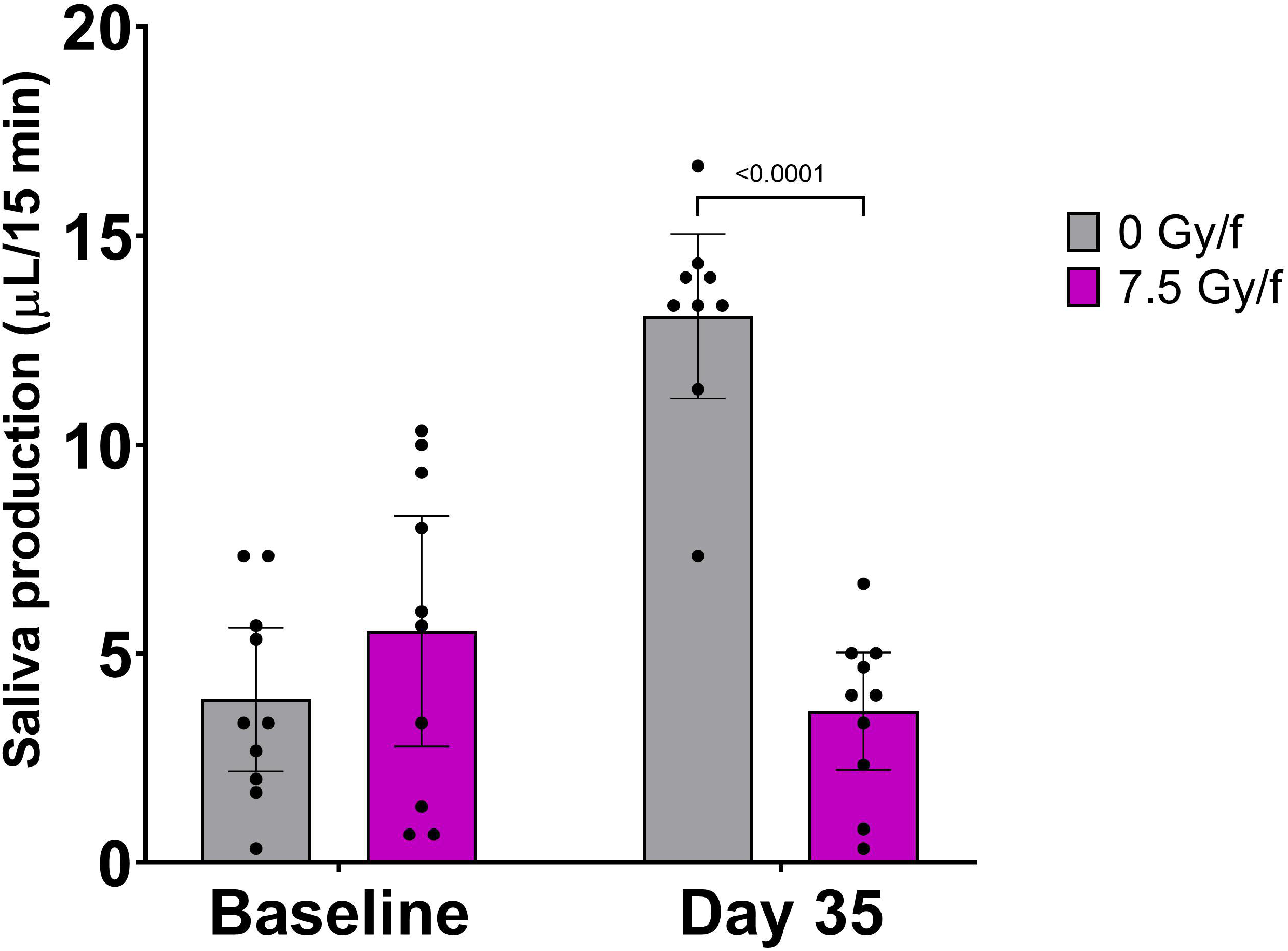
Significantly lower saliva production was found in irradiated mice compared to controls at day 35 after exposure to fractionated irradiation. Saliva production was measured as µL/15 min in 7.5 Gy/f and controls. Data is represented as mean ± 95% CI. Each black dot represents an animal.

## Discussion

In the current work, we show early radiation-induced effects using our preclinical model in which mice are exposed to fractionated radiation in the head and neck region. In the present study, we have tested doses per fraction ranging from 3-8.5 Gy x 10 (30-85 Gy total dose) and have found that for the current fractionation scheme and irradiation setup, the optimal dose to obtain clinically relevant normal tissue response is 7.5 Gy/f. After exposure to 7.5 Gy/f mice experienced radiation dermatitis in the skin of the lip and upper chest, oral mucositis in the ventral tongue, acinar vacuolization in the parotid glands and significantly reduced saliva production compared to control mice.

Early radiation-induced effects in the H&N region depend on various parameters including type of radiation, dose, radiation field configuration, and irradiation protocol (single dose (SD) or fractionation). In preclinical research, the selection of different rodent strains for such studies also influences the results due to strain-specific variations in response to RT-induced damage such as oral mucositis (20) and morphological and functional changes of the salivary glands (21). Radiation-induced side effects of the H&N have previously been studied in hamsters (22, 23), rats (24-26), and mice (7, 8, 10, 13, 27). Due to the lack of assay reagents available for hamsters, and that it is easier to keep a larger number of mice than hamsters or rats, mice were chosen for the present preclinical model. It is important to use a model that can cover both tumor and normal tissue effects in order to estimate the therapeutic ratio (tumor vs normal tissue effects) and to measure the effect of mitigators on late effects. We therefore selected C57BL/6 mice as this mouse strain has a large panel of syngeneic tumors. Compared to other mouse strains, C57BL/6 is relatively radioresistant (28-30) and is therefore a suitable strain for studying both early and late radiation-induced effects in the same model, as the mice will tolerate high doses of radiation. It is also the most widespread substrain used for studying genetically engineered mice (31) and has been recommended for studies with radioprotectors and mitigators (32). It is well known that inbred mice are roughly half as sensitive to radiation as humans (30). However, relevant exposures and regimens can be found by comparing effects seen in mice with effects observed in humans.

With the selected mouse strain, we established a setup including clinically applicable radiation fields with added imaging and MC simulations to estimate local radiation doses to relevant tissues. Compared to 180 kV X-rays, the simulations showed that 100 kV X-rays produced a more relevant, heterogeneous dose distribution, similar to clinical radiotherapy focusing on one part of the H&N region. In addition, fractionated irradiation was employed because of higher clinical relevance compared to SD. Dose fractionation in preclinical studies has not been extensively reported in the literature. Moreover, previous studies have either focused on early or late effects, while we believe that studying both early and late phases of the different tissue reactions could show the possible relationship between these effects and the influence of systemic processes (e.g., cytokine expression). In the present study we only focused on early radiation-induced effects, however the preclinical model presented here will be included in experiments with longer follow-up periods to study late radiation-induced effects.

In our study, the rate and severity of the skin toxicity was dependent on radiation dose, as evident from dose response curves and LD_50_ values. The RTOG-based skin toxicity scoring table employed was not sufficiently sensitive to distinguish between all the tested doses per fraction. However, it could distinguish between 3 different groups: low dose (3 and 4.4 Gy/f), intermediate dose (5-6.5 Gy/f), and high dose (7.5 and 8.5 Gy/f). The macroscopic and microscopic changes seen in the skin of the lower lip were compatible with radiation dermatitis reported in mouse skin after SD (33) and fractionated irradiation with fewer fractions than we used (7, 14, 34, 35). Even though the irradiation protocol and doses differ between the present work and earlier studies, the histological changes in the skin are similar. This might indicate that radiation-induced dermatitis occurs independent of irradiation protocol and dose, as long as the dose is above a certain threshold level.

Oral mucositis was observed macroscopically and histologically in the ventral tongue of mice exposed to 7.5 and 8.5 Gy/f. This is similar to what has been reported 8 days after SD irradiation of 18-25 Gy (8, 36), 10 days after fractionated irradiation given as 10 Gy/f over 3 days (30 Gy in total dose) (7) and as 3 Gy/f over 5 days (15 Gy in total dose) (37). However, not all studies on oral mucositis in rodents observe oral mucosal ulcers. In other studies using SD irradiation, the authors state that rodents do not develop oral mucositis in the same manner as humans, but progress towards weight loss and death before ulcerative lesions appear (9, 10). This is in contrast to the ulcerative lesions we observe in the ventral tongue together with weight loss in mice after exposure to 7.5 Gy/f. Both these effects were non-lethal and temporary for the given dose level.

Saliva production was significantly lower in irradiated mice (7.5 Gy/f) compared to controls at day 35, which is consistent with previous reports after SD irradiation in mice (38-41) and fractionated irradiation in mice and rats (6, 25). Interestingly, at day 35 saliva production in controls increased significantly from baseline, which might indicate that saliva production increases with age in normal salivary glands. Indeed, it has been shown that C57BL/6 mice increase their saliva production as they age from 10-weeks-old to 30-weeks-old (42), which is in line with our results. In irradiated salivary glands however, the saliva production at day 35 was slightly lower compared to baseline, implying that salivary gland function was compromised. The observed saliva levels in the present experiments varied considerably between animals and time points and decreased with increasing dose (average CV of 59 %). Additionally, we observed acinar vacuolization in the parotid glands in mice exposed to 8.5 Gy/f, which is consistent with previous reports after SD irradiation in mice (5, 6, 43). The damages in the parotid glands may be parts of the explanation of the reduced saliva production in irradiated mice compared to controls.

In conclusion, the proposed preclinical model allows for the study of early radiation-induced effects in the H&N region. We observed both macroscopic and microscopic changes in skin, oral mucosa of tongue and lip, and salivary glands as well as a significantly reduced saliva production in RT compared to controls. The present preclinical model is an optimal compromise with respect to ease of use and possibilities for short term studies and for future long term follow-up with tissue collection and liquid sampling that can allow for cytokine analysis in blood and/or saliva. Regarding the X-ray energies used, our Monte Carlo simulations show that 100 kV X-rays provide a more relevant and heterogeneous dose distribution compared to 180 kV, which will provide a better basis for future studies comparing side effects from X-rays and ions such as protons. In the present study, we have tested doses per fraction ranging from 3-8.5 Gy and have found that for the current fractionation scheme and irradiation setup, the optimal dose is 7.5 Gy/f. This dose is well tolerated in this mouse strain and results in similar early radiation-induced effects as in H&N cancer patients.

## Supporting information

https://drive.google.com/drive/folders/1ac0VKx3YzlATLM_NYqoLfThwNoXDlCrj?usp=sharing

