## Supplementary material for "A preclinical model to investigate normal tissue damage following fractionated radiotherapy to the head and neck": https://drive.google.com/drive/folders/1ac0VKx3YzlATLM_NYqoLfThwNoXDlCrj?usp=sharing

### Supplementary file

#### Supplementary Materials and Methods

***Monte Carlo simulations***

DICOM computed tomography (CT) images of one euthanized male mouse (10-11 weeks of age) with isotropic voxel sizes of 0.02 × 0.02 × 0.02 $mm^3$ were imported into Matlab R2020b (MathWorks, Natick, MA, USA) and co-registered to diagnostic DICOM magnetic resonance (MR) images of voxel sizes 0.12 × 0.12 × 0.70 $mm^3$ in the sagittal, coronal and axial plane of the animal, respectively, using an in-house developed software. Registration-based interpolation was performed in order to spatially align the axial CT stack to the axial MR images (fixed, reference) when equivalent posture and positioning of the mouse between the CT-MR image scanning acquisitions was accomplished. The interpolated CT images were subsequently imported into FLUKA 4-1.1 [1-3], where the geometrical scheme of the irradiation setup was defined using DICOM tools in Flair v3.1-13 (FLUKA Advanced Interface) [4]. Herewith, FLUKA Monte Carlo (MC) simulations allowed for exploration of the tissue dose distribution following different applied X-ray voltages. The absorbed dose distributions were simulated for 100 kV and 180 kV X-ray spectra attenuated by 2.0 mm Al and 0.3 mm Cu filters, respectively. For both energies, a rectangular irradiation field of size 1.5 × 0.75 $cm^2$ was focused sagittally to the neck region of the mouse from a 52.0 cm source to sample distance (SSD), providing salivary gland area coverage and other normal tissue structures.

For the voxelized CT-based geometry, the FLUKA MC system was calibrated to match tissue mass densities and stopping powers with correlating CT numbers by a published stoichiometric calibration procedure [5]. Moreover, the MC simulations were run using PRECISIOn defaults to guarantee reliable accuracy without increasing the simulation time significantly, where the transport and production cutoff of photons and electrons was set to 1 keV. The treatment field was simulated for $5 \times 10^7$ primary X-ray photons. Furthermore, dose scoring was performed using the USRBIN card in FLUKA. The dose quantities were scored on a voxel-by-voxel basis using the grid specified by the aforementioned MR images, wherefore the scoring region was defined to encompass all relevant volumes.

***Immunohistochemistry***

For immunohistochemistry, tissue sections of 4 µm were cut (Leica RM2155 microtome) and placed on microscopy glass slides (Superfrost Plus, Thermo Fisher). Following deparaffination and hydration, heat-induced antigen retrieval was performed using citraconic anhydride pH 7.4, for 15 minutes at 100 ºC. Primary antibodies against CD138/Syndecan-1 (Rat IgG2a, 1:450), CD3 (Rat IgG1, 1:450), Vimentin (clone EPR3776, Rabbit IgG, 1:500) and F4/80 (clone BM8, Rat IgG2a, #123101, BioLegend) were used. Before incubation with primary antibodies, blocking was performed for 60 minutes at room temperature using Normal Rabbit Serum 5 % (CD138, CD3, F4/80) and Normal Goat Serum 5 % (Vimentin). Next, sections were incubated with primary antibodies over night at 4 ºC. Negative control was PBS instead of primary antibody. Positive controls were sections of mouse spleen (CD3, CD138, F4/80), plasmacytoma (CD3, CD138) and blood vessels (Vimentin). Furthermore, the sections were washed in PBS for 2 x 10 minutes and incubated with secondary antibodies for 40 minutes at room temperature. Secondary antibodies used were Rabbit-a-Rat IgG (CD3, CD138, F4/80) and Goat-a-Rabbit IgG (Vimentin). Next, the slides were washed with PBS 2 x 10 minutes, incubated with biotinylated streptavidin ($ABC^{HRP}$) for 30 minutes at room temperature, incubated with DAB for 10 minutes at room temperature and washed with PBS. Finally, the slides were stained with Hematoxylin for 30 seconds, dehydrated and mounted with synthetic resin.

Histological images were acquired using a Nikon DS-Ri1 camera with a CFI Plan Fluor 20x objective (NA 0.5).

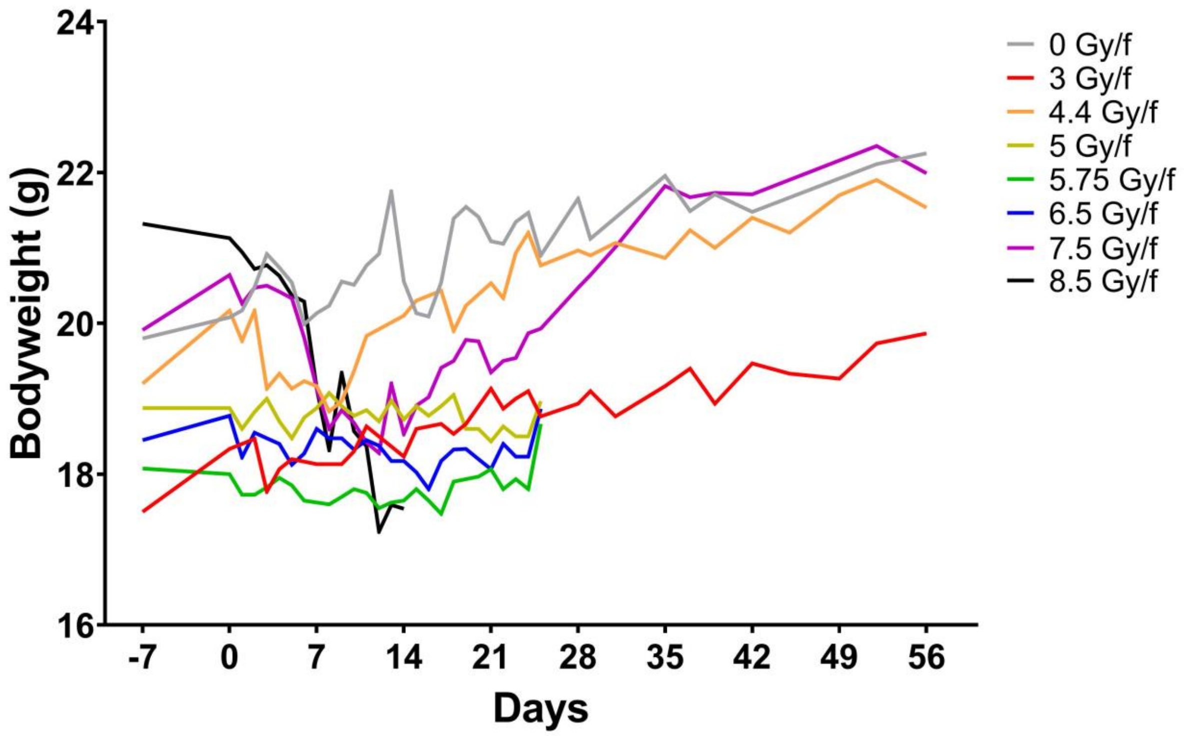


Supplementary figure 1. Bodyweight loss greater than 10 % was only seen in the mice exposed to 7.5 and 8.5 Gy/f (black and purple group). Mice receiving 7.5 Gy/f increased in weight and recovered around day 14, while mice receiving 8.5 Gy/f did not recover and was euthanised at day 14. The graphs show mean bodyweight in each group. The number of animals used in each group varied as following 0 Gy/f (n = 9), 3 Gy/f (n = 3), 4.4 Gy/f (n = 3), 5 Gy/f (n = 4), 5.75 Gy/f (n = 4), 6.5 Gy/f (n = 4), 7.5 Gy/f (n = 10), 8.5 Gy/f (n = 10). The black vertical line on the X-axis represent each day of radiation, giving 2 fractions per day.

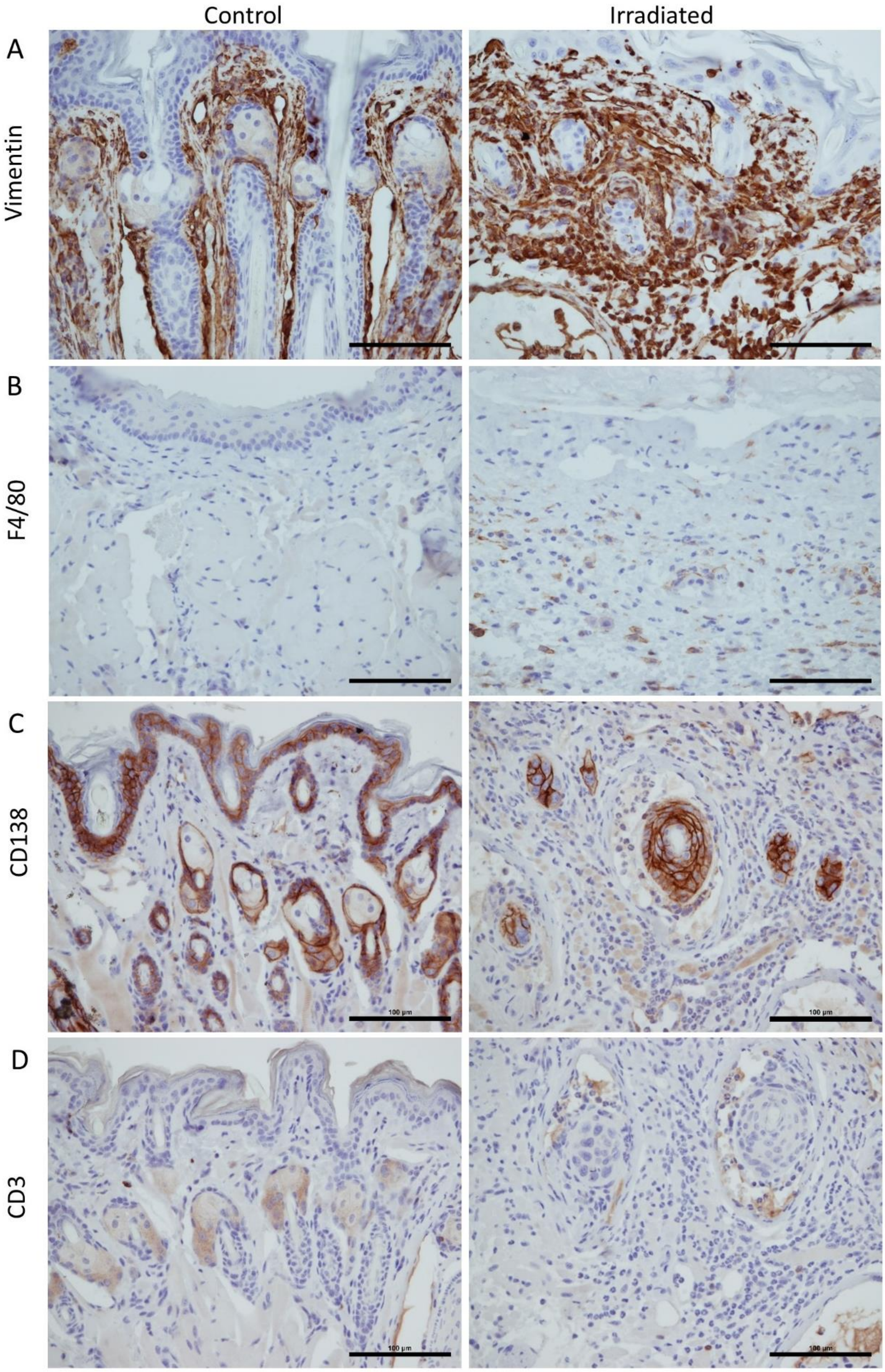


Supplementary figure 2. Immunohistochemically labelled sections of the lower lip in mice after exposure to fractionated irradiation. (A-B) Irradiated sections showed an increase in Vimentin$^{+}$ cells and F4/80$^{+}$ cells. Fibroblasts as well as neutrophils and lymphocytes are positive for Vimentin, while macrophages in mice are F4/80$^{+}$. (C) CD138 labeled epithelial cells, but showed no increase of plasma cells in the irradiated sections. (D) CD3 labelled the sebaceous glands, but showed no obvious increase of T cells in the irradiated sections. Scale bar is 100 µm.

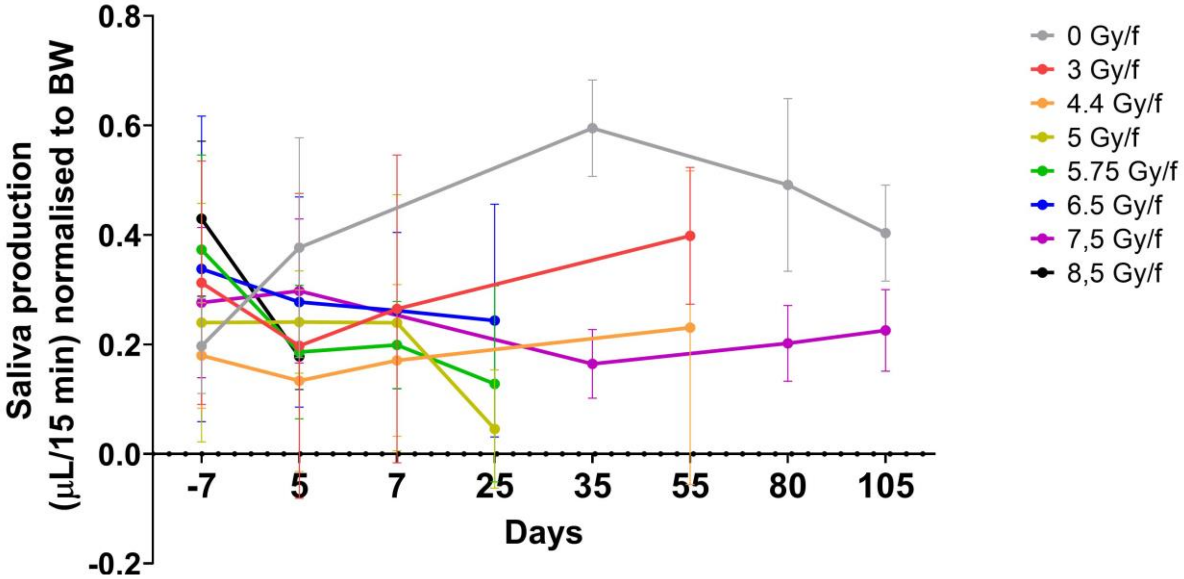

Supplementary figure 3. Saliva production measured as µL saliva/15 minutes normalised to bodyweight. Data is represented as mean ± 95% CI. The number of animals used in each group varied as following 0 Gy/f (n = 9), 3 Gy/f (n = 3), 4.4 Gy/f (n = 3), 5 Gy/f (n = 4), 5.75 Gy/f (n = 4), 6.5 Gy/f (n = 4), 7.5 Gy/f (n = 10), 8.5 Gy/f (n = 10).

***References***


1. Battistoni, G., et al., *Overview of the FLUKA code.* Annals of Nuclear Energy, 2015. **82**: p. 10-18.
2. Böhlen, T.T., et al., *The FLUKA Code: Developments and Challenges for High Energy and Medical Applications.* Nuclear Data Sheets, 2014. **120**: p. 211-214.
3. Battistoni, G., et al., *The FLUKA Code: An Accurate Simulation Tool for Particle Therapy.* Frontiers in oncology, 2016. **6**: p. 116-116.
4. Vlachoudis, V., *Flair: A powerful but user friendly graphical interface for FLUKA.* 2009, United States: American Nuclear Society - ANS.
5. Schneider, W., T. Bortfeld, and W. Schlegel, *Correlation between CT numbers and tissue parameters needed for Monte Carlo simulations of clinical dose distributions.* Phys Med Biol, 2000. **45**(2): p. 459-78.
